# A spike-sorting method based on hierarchical clustering to discriminate extracellularly recorded simple spikes and complex spikes from cerebellar Purkinje cells

**DOI:** 10.1101/2021.09.30.462526

**Authors:** Takayuki Michikawa, Keisuke Isobe, Shigeyoshi Itohara

## Abstract

**Background:** In the cerebellar cortex, Purkinje cells are the only output neurons and exhibit two types of discharge. Most Purkinje cell discharges are simple spikes, which are commonly appearing action potentials exhibiting a rich variety of firing patterns with a rate of up to 400 Hz. More infrequent discharges are complex spikes, which consist of a short burst of impulses accompanied by a massive increase in dendritic Ca^2+^ with a firing rate of around 1 Hz. The discrimination of these spikes in extracellular single-unit recordings is not always straightforward, as their waveforms vary depending on recording conditions and intrinsic fluctuations.

**New Method:** To discriminate complex spikes from simple spikes in the extracellular single-unit data, we developed a semiautomatic spike-sorting method based on divisive hierarchical clustering.

**Results:** Quantitative evaluation using parallel *in vivo* two-photon Ca^2+^ imaging of Purkinje cell dendrites indicated that 96.6% of the complex spikes were detected using our spike-sorting method from extracellular single-unit recordings obtained from anesthetized mice.

**Comparison with Existing Method(s):** No reports have conducted a quantitative evaluation of spike-sorting algorithms used for the classification of extracellular spikes recorded from cerebellar Purkinje cells.

**Conclusions:** Our method could be expected to contribute to research in information processing in the cerebellar cortex and the development of a fully automatic spike-sorting algorithm by providing ground-truth data useful for deep learning.

**Highlights:**

- A spike-sorting algorithm on hierarchical clustering was developed.
- It was applied to extracellular recordings from cerebellar Purkinje cells.
- Complex and simple spikes were discriminated based on their waveforms.
- Spike-sorting performance was quantified using *in vivo* Ca^2+^ imaging data.
- The algorithm isolated over 96.6% of complex spikes.

## 1. Introduction

Purkinje cells are the only output neurons in the cerebellar cortex, and each receives two different types of excitatory inputs: one consists of thousands of weak inputs from parallel fibers of granule cells, and the other consists of a strong input from a single climbing fiber of an inferior olive neuron (Eccles et al., 1967) (Fig. 1A). A single Purkinje cell shows two types of discharge: a simple spike (SS) and a complex spike (CS) (Thach, 1967) (Fig. 1A). The SS activity pattern exhibits a rich variety (De Zeeuw et al., 2011; Xiao et al., 2014), including tonic firing with a firing frequency of up to 400 Hz (Monsivais et al., 2005) and bistable firing in which tonic firing in the up-state alternates with quiescent periods in the down-state (Loewenstein et al., 2005; Schonewille et al., 2006; Yartsev et al., 2009). The SS firing pattern is modulated by various factors, such as the intrinsic activity of Purkinje cells, excitatory inputs from parallel fibers, and inhibitory inputs from molecular layer interneurons (De Zeeuw et al., 2011). A CS is a short burst of pulses consisting of an initial spike followed by several spikelets (Granit and Phillips, 1956), and is evoked by the activation of a climbing fiber (Eccles et al., 1966). Single action potentials in olive neurons are translated in the burst of climbing fiber spikes, and the number of spikes in the burst depends on the phase of the subthreshold membrane potential oscillations in olive neurons (Mathy et al., 2009). The number of spikes within the presynaptic climbing fiber burst can influence both the number of spikelets and the duration of the postsynaptic CSs generated in the Purkinje cell axon (Davie et al., 2008; Mathy et al., 2009). Purkinje cell plasticity and cerebellar motor learning are graded depending on CS duration (Mathy et al., 2009; Rasmussen et al., 2013; Yang and Lisberger, 2014a). Therefore, in addition to discriminating CSs from SSs accurately in extracellular recordings, classifying CSs depending on their duration is critical for understanding information processing in the cerebellar cortex.

**Fig. 1.**
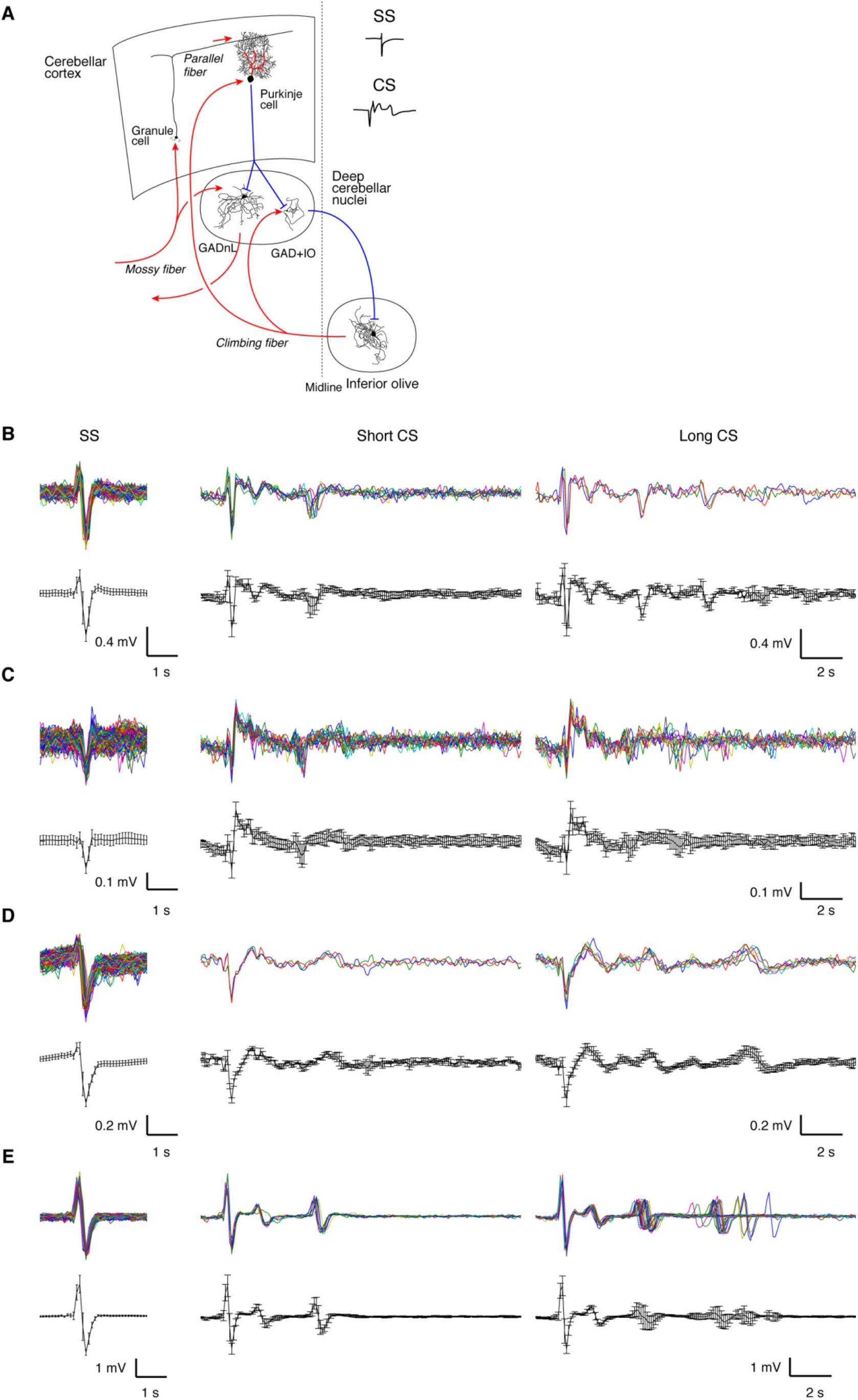
Variability of extracellular SS and CS waveforms recorded from cerebellar Purkinje cells in anesthetized mice. (A) A schematic drawing of the main circuits in the cerebellum. Excitatory and inhibitory inputs are shown in red and blue, respectively. GADnL: glutamic acid decarboxylase (GAD)-negative non-GABAergic large projection neuron (Uusisaari and Knopfel, 2011). GAD+IO: GAD-positive GABAergic neuron projecting to the inferior olive (Uusisaari and Knopfel, 2011). SSs are produced endogenously in PCs, and their firing patterns are modulated by excitatory synaptic inputs from parallel fibers that relay various inputs delivered from mossy fibers that arise from throughout the brain and spinal cord via several brain stem and pontine nuclei. A CS is evoked by a climbing fiber input originating from the contralateral inferior olive. (B–E) Extracellular spike waveforms recorded from four different Purkinje cells. Typical SS waveforms and two (short and long) CS waveforms are shown for each cell. Individual waveforms are superimposed in the upper panels, and their average waveforms are shown in the bottom panels. Error bars: standard deviation.

From a practical point of view, it is not always straightforward to separate CSs from SSs in extracellular single-unit Purkinje cell records because this involves a number of difficulties: (1) CSs and SSs may have similar waveforms, (2) CS waveforms vary from cell to cell, (3) CS waveforms may change over time within a single record, and (4) the number and timing of spikelets in CSs may vary substantially (Rasmussen et al., 2014). Off-line spike sorting of Purkinje cell firing can be performed using a variety of methods or algorithms, including (1) principal component analysis (Kitazawa et al., 1998), (2) window discriminators (Catz et al., 2005; Yang and Lisberger, 2014b), (3) template-matching algorithms (Wise et al., 2010), (4) thresholding of square or cube waveforms of filtered extracellular signals (Rasmussen et al., 2014), (5) comparison with the average waveform of hand-selected CSs (de Solages et al., 2008; Warnaar et al., 2015), and (6) full visual selection (Bosman et al., 2010; Soetedjo et al., 2008). Some investigators select only extracellular records with a sufficient signal-to-noise ratio beforehand to discriminate between CSs and SSs reliably. However, to our knowledge, no reports have conducted a quantitative evaluation of spike-sorting algorithms used for the classification of extracellular spikes recorded from cerebellar Purkinje cells *in vivo*.

Hierarchical clustering is a cluster analysis method that seeks to construct a hierarchy of observation clusters. There are two strategies to perform hierarchical clustering: an agglomerative approach and a divisive approach. The former is a bottom-up approach that starts in its own cluster, and pairs of clusters are merged as one moves up the hierarchy. The latter is a top-down approach that starts in a single cluster, and splits are performed recursively as one moves down the hierarchy. The clustering depends on a measure of distance between pairs of observations and a linkage criterion that determines the dissimilarity of sets as a function of the pairwise distances of observations in sets. The results of hierarchical clustering are usually presented in a dendrogram in which the distances between clusters are represented graphically.

In the present study, we applied divisive hierarchical clustering analysis to discriminate CSs from SSs in extracellular single-unit data recorded from cerebellar Purkinje cells in anesthetized mice. Hierarchical clustering analysis is also used to classify CSs according to spikelet shape. Since the firing of CSs is accompanied by an increase of Ca^2+^ in postsynaptic Purkinje cell dendrites (Miyakawa et al., 1992), we simultaneously measured dendritic Ca^2+^ using *in vivo* two-photon Ca^2+^ imaging to evaluate the performance of the spike-sorting algorithm. The results provide a quantitative measure of the rate of correctly sorted CSs using the algorithm.

## 2. Materials and Methods

### 2.1. Animal preparation

All experimental procedures were performed in accordance with the guidelines of the Animal Experiment Committee of the RIKEN Center for Brain Science. ICR mice were anesthetized with a mixture of fentanyl (0.05 mg/kg), midazolam (5.0 mg/kg), and medetomidin (0.5 mg/kg), according to a method described previously (Mrsic-Flogel et al., 2007). Anesthesia was maintained by reinjecting one third of the initial dose at appropriate intervals.

### 2.2. Expression of Ca^2+^ indicator proteins in cerebellar Purkinje cells

Recombinant adenovirus carrying yellow cameleon 2.60 (YC2.60) (Nagai et al., 2004) was injected into the lateral ventricle of mice embryos at embryonic day 11, as described previously (Yamada et al., 2011). Mice expressing YC2.60 were identified during postnatal days 1–3 by viewing their transcranial fluorescence.

### 2.3. Surgery

In acute experiments, a custom-made stainless steel headplate was attached to the mouse skull using surgical cyanoacrylate and dental cement (Super-bond C&B, Sun Medical, Shiga, Japan) under anesthesia on the day of measurement. The cerebellar surface was covered by 2% agarose and 0.17-mm thick cover glass (Matsunami Glass, Osaka, Japan) after removing a piece of the skull and dura mater.

### 2.4. Electrophysiology and *in vivo* Ca^2+^ imaging

Extracellular single-unit recording and dendritic Ca^2+^ imaging of cerebellar Purkinje cells in anesthetized head-fixed mice were performed on postnatal days 30–220. Extracellular single unit recording was carried out using borosilicate glass pipettes (1–3 MΩ) filled with extracellular solution containing 150 mM NaCl, 2.5 mM KCl, 10 mM HEPES, 2 mM CaCl_2_, 1 mM MgCl_2_, and 0.1 mg/ml dextran AlexaFluor 568 (pH 7.3, ~300 Osm/L) by targeting to a YC2.60-expressing Purkinje cell soma visualized with an upright two-photon laser-scanning microscope (LSM710MP; Carl Zeiss, Jena, Germany). Extracellular signals were amplified 100 times and acquired at 100 kHz using a microelectrode current and voltage clamp amplifier (MultiClamp 700B; Molecular Devices, San Jose, CA) and a low-noise digitizer (Digidata 1440A; Molecular Devices) controlled by pClamp 10 software (Molecular Devices). Fluorescent signals of YC2.60 were simultaneously acquired in frame-scan mode (256 × 256 pixels, 2–24 Hz) using a 20’ water-immersion objective (W Plan-Apochromat 20×/1.0 DIC D×0.17 M27 70 mm). Image acquisition faster than 4 Hz was carried out using a line-step mode equipped in ZEN2010 software (Carl Zeiss) with a pair of Galvano scanners. A Ti:sapphire laser (Chameleon Ultra, Coherent, Santa Clara, CA) for excitation was tuned to 820 nm. Emitted fluorescence was short-pass filtered (630 nm), split with a dichroic mirror (510 nm), band-pass filtered (460–500 nm and 525–560 nm for cyan and yellow fluorescence, respectively), and detected with GaAsP detectors. The electrophysiological recordings and two-photon imaging were synchronized by a trigger pulse generated using ZEN2010 image acquisition software.

### 2.5. Data analysis

Electrophysiological and fluorescence signals were analyzed using custom-made software written in MATLAB. A detailed description of the spike-sorting software, *spike_hiclus,* is provided in the supplemental material (Figs. S1–S6). The image data analysis method is described elsewhere (Michikawa, T., et al., 2020).

## 3. Results

### 3.1. Variability in extracellular SS and CS waveforms

The SS and CS waveforms in extracellular single-unit recordings of four different Purkinje cells are shown in Fig. 1. SSs exhibited biphasic positive-negative potentials (Fig. 1B,D,E), monophasic negative potentials (Fig. 1C), and a mix of both (see below). CS waveforms also showed cell-to-cell variability, and their initial peaks (positive–negative potentials or negative potentials) were similar to each cell’s SS waveform. Spikelets of CSs followed by the initial peak also exhibited remarkable cell-to-cell variability. The amplitude of the second positive potential was comparable to the peak height of the first positive potential in the cell shown in Fig. 1B, while the amplitude of the second positive potential was larger than the peak height of the first positive potential in the cell shown in Fig. 1C. In the cell shown in Fig. 1D, the first positive potential was very small, and the peak of the second positive potential was delayed approximately 1 ms after the peak of the first negative potential. In addition to the variability in spike waveforms, the number of spikelets in CSs was variable, even in data recorded from the same cells. Typical examples of short CSs lasting approximately 5 ms and long CSs lasting approximately 10 ms are shown for each cell in Fig. 1. Apart from the variability, completely different waveforms were occasionally recorded. The voltage signals exhibited relatively large biphasic positive–negative potentials (Fig. 1E), similar to those obtained in juxtacellular recordings (Pinault, 1996). The initial potentials of CSs in these records were almost the same as the SS waveforms, with two or three additional well-separated, small, blunt biphasic spikes followed by variable intervals. The spike-sorting algorithm needs to distinguish SSs and CSs over these variabilities correctly.

### 3.2. The spike-sorting algorithm based on hierarchical clustering

A flow diagram of the spike-sorting algorithm based on hierarchical clustering is shown in Fig. 2. The algorithm has four main stages: (1) event detection via amplitude thresholding; (2) hierarchical clustering to classify SSs, CSs, and noise; (3) manual correction of both SSs and CSs; and (4) hierarchical clustering to classify CSs depending on the number or shape of spikelets. In this study, extracellular single-unit recordings were made using simultaneous two-photon Ca^2+^ imaging from the same Purkinje cell. By using the dendritic Ca^2+^ increase as the index of occurrence of a CS, we could evaluate the performance of the spike-sorting algorithm, particularly for discriminating CSs. By comparing the events classified as CSs and the two-photon Ca^2+^ imaging data, as shown in the broken enclosure in Fig. 2, we could quantify the correct sorting rate of CSs and isolate false and missed events if present. If there were missed events, we attempted to identify the corresponding CSs from the events classified as SSs. If the corresponding events did not exist in the SS clusters, the novel events were selected as CSs from the original extracellular single-unit traces.

**Fig. 2.**
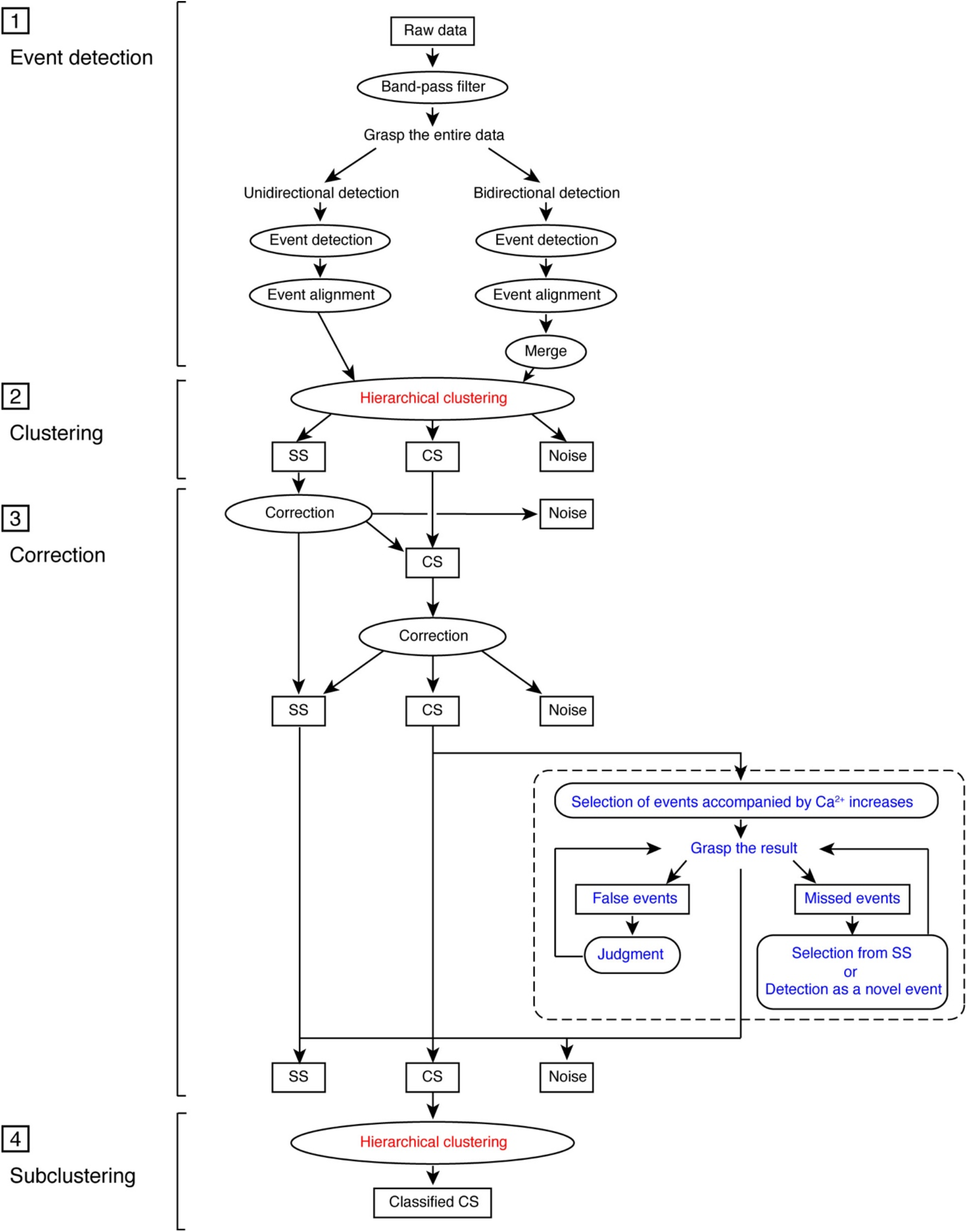
Flow diagram of the spike-sorting algorithm using hierarchical clustering. The computational processes are illustrated in ellipses. The processes with two-photon Ca^2+^ imaging data are shown in dashed enclosure.

### 3.3. Hierarchical clustering of events

The results of hierarchical clustering of the data including monophasic SSs are shown in Fig. 3. After visual inspection of the raw traces, a single negative threshold was chosen for event detection. A threshold of –5σ_n_ (Quiroga et al., 2004) was chosen, where σ_n_ is an estimate of the standard deviation of the background noise, computed as follows:

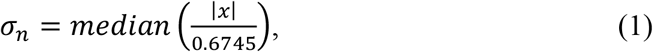

where *x* is the bandpass-filtered single-unit signal. The event time was aligned with the time of the minimal voltage signal within –1.5 ms to +0.5 ms of the threshold crossing time. The waveforms of each event were constructed from –1.5 ms to +2 ms of the event time and applied to hierarchical clustering. Ward’s linkage was used as the algorithm for computing distances between clusters *r* and *s*, *d(r,s)*, as follows:

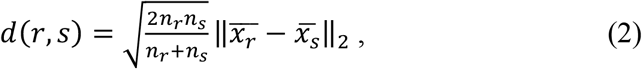

where || ||_2_ is the Euclidean distance, 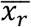, and 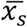 are the centroids of clusters *r* and *s*, and *n_r_* and *n_s_* are the numbers of elements in clusters *r* and *s*, respectively.

**Fig. 3.**
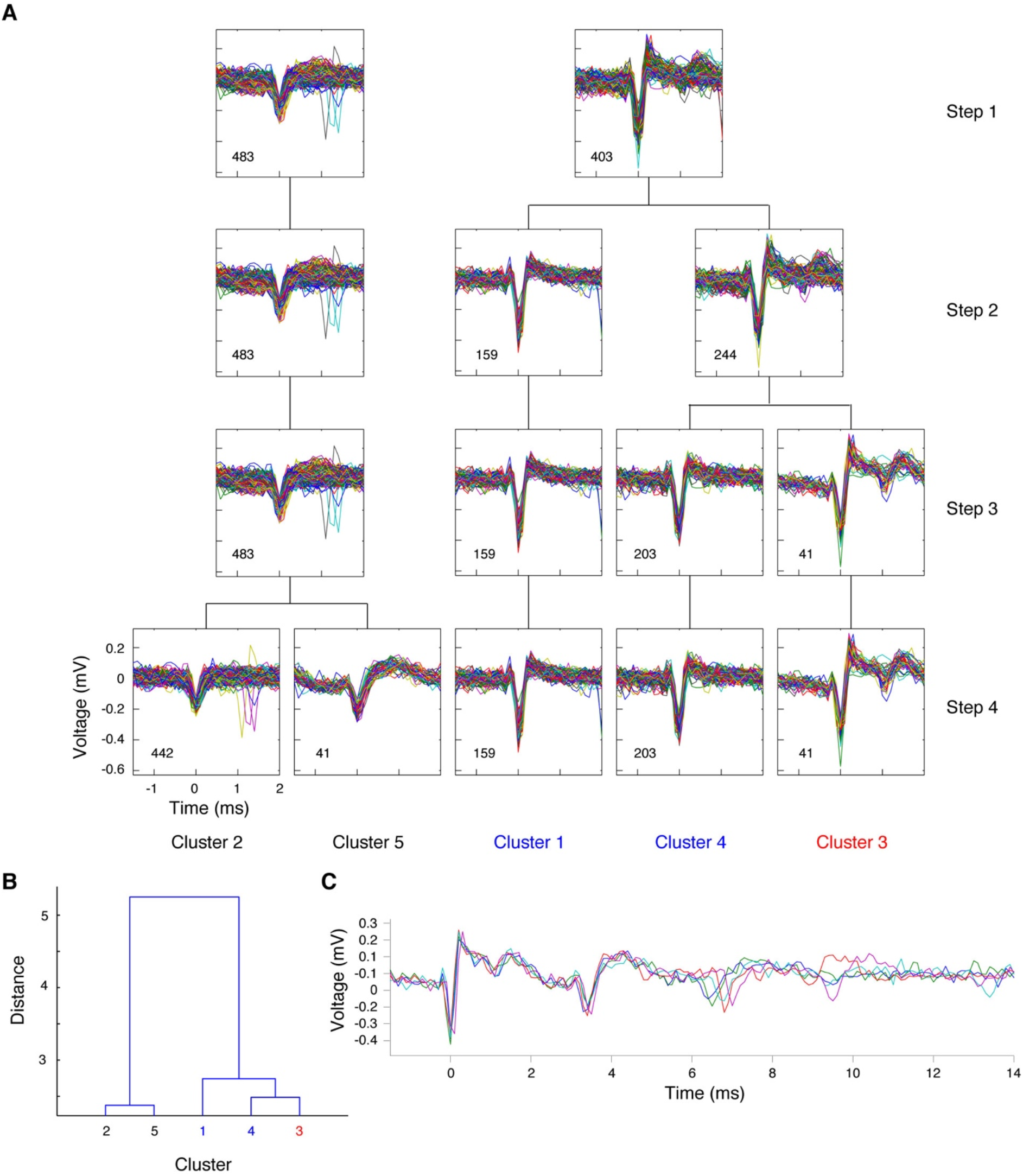
Spike sorting of data including monophasic SSs. (A) Results of hierarchical clustering that classified spikes detected in a 100-s recording into five clusters. The waveforms of individual spikes are superimposed with different colors in each plot. The number of spikes is shown in each panel. The cluster classified as CSs is shown in red (cluster 3), and the clusters classified as SSs are shown in blue (clusters 1 and 4). (B) Dendrogram of hierarchical clustering. (C) The waveforms classified as CSs are superimposed (n = 5).

The results of the four-step hierarchical clustering are shown in Fig. 3A. In the first step, we split the 886 events isolated from the 100-s record into two clusters composed of 483 and 403 events, respectively (Fig. 3A). In the second step, we split the cluster composed of 403 events in the right branch into two clusters of 159 and 244 events, respectively. In the third step, we split the cluster composed of 244 events in the right branch into two clusters of 203 and 41 events, respectively. In the fourth step, we split the cluster composed of 483 events in the left branch into two clusters of 442 and 41 events, respectively. At this stage, hierarchical clustering was terminated by visual inspection. A dendrogram of the hierarchical clustering (Fig. 3B) indicated that the left and right branches were remarkably different. The cluster composed of 41 events of the right branch, cluster 3, was defined as CSs and the clusters composed of 159 events (cluster 1) and 203 events (cluster 4) in the right branch were defined as SSs.

The hierarchical clustering result (Fig. 3A) demonstrates the difficulty involved in the spike sorting of extracellular single-unit recordings obtained from Purkinje cells. We constructed spike waveforms with a duration of 3.5 ms (event time from –1.5 ms to +2 ms), since longer durations prevent accurate spike sorting when a cell fires at a high frequency, such as > 200 Hz, and shorter durations tend to fail for discriminating CSs from SSs. Using this duration, CS spikelets were isolated as separate events. The cluster composed of 41 events of the left branch, cluster 5, might have corresponded to spikelets at approximately 3 ms after the onset of CS (Fig. 3C). Therefore, we excluded cluster 5 from CSs and SSs. A cluster composed of 442 events in the left branch, cluster 2, may also have contained spikelets appearing approximately 6 ms after CS onset (Fig. 3C). Since clusters 1 and 4 in the fourth step were classified as the same cluster in the first step, these spikes could have originated from the same Purkinje cells, and cluster 1 corresponded to SSs.

The results of hierarchical clustering of the data including biphasic SSs are shown in Fig. 4. We detected events with a uniform threshold set to –5σ_n_. After seven steps of hierarchical clustering, CSs and CS spikelets were isolated as clusters 6 and 8, respectively. SSs were divided into six clusters (clusters 2–4 and 7 in the left branch and clusters 1 and 5 in the right branch).

**Fig. 4.**
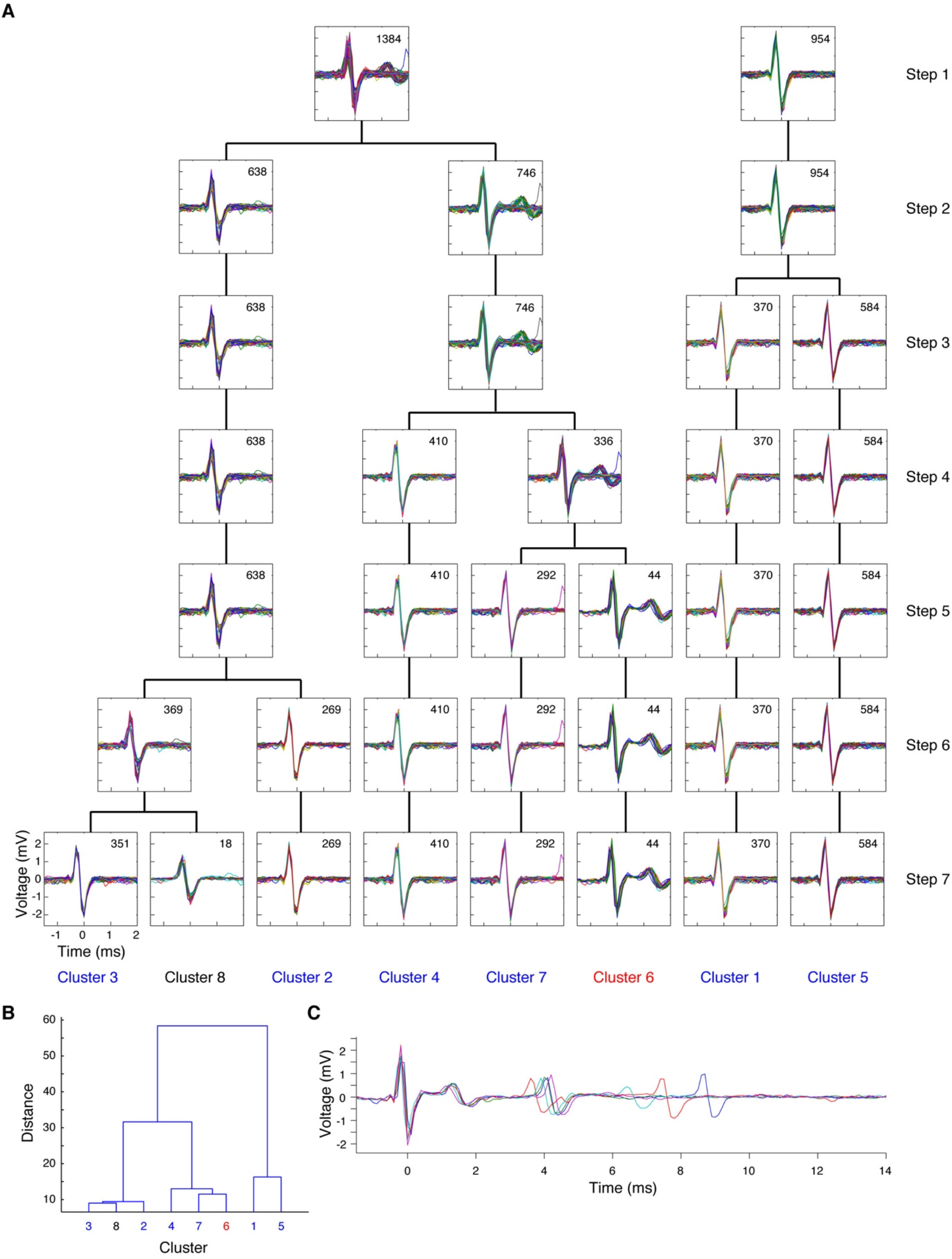
Spike sorting of data including biphasic SSs. (A) Results of hierarchical clustering that classified spikes detected in a 100-s recording into eight clusters. The waveforms of individual spikes are superimposed with different colors in each plot. The number of spikes is shown in each panel. The cluster classified as CSs is shown in red (cluster 6) and the clusters classified as SSs are shown in blue (clusters 1–5 and 7). (B) Dendrogram of hierarchical clustering. (C) The waveforms classified as CS are superimposed (n = 5).

The results of hierarchical clustering of the data containing CSs without initial negative potentials are shown in Fig. 5A. In this case, events were detected with both positive and negative thresholds set to +5σ_n_ and –5σ_n_, respectively. For events detected with the negative threshold, the event time was aligned to the minimum peak time within –1.5 ms to 0.5 ms of the threshold crossing time, whereas for events detected with the positive threshold, the event time was aligned to the maximal peak time within –1.5 ms to 0.5 ms of the threshold crossing time. Both events were merged before hierarchical clustering. To avoid multiple selections of single events in the merged data, we adapted only those events that were apart by at least 1.5 ms from the preceding event. For the first step of hierarchical clustering, we split the merged 1,413 events into two clusters of 332 and 1,081 events, respectively. The cluster composed of 332 events was detected with the positive threshold, since the maximum was at time 0 (Fig. 5A). The other cluster composed of 1,081 events was detected with the negative threshold because the minimal was at time 0 (Fig. 5A). After 10 steps of hierarchical clustering, CSs were sorted in clusters 6, 9, and 11. The spikelets that appeared approximately 4 ms after the onset of CSs (Fig. 5C) were sorted in cluster 7 (Fig. 5A). SS classification was difficult in this case because there were many different waveforms in both the left and right branches. Biphasic spikes sorted in cluster 8 and 10 were most likely to be SSs since the initial positive peaks resembled the initial peaks of the CSs. A large number of monophasic spikes (clusters 3 and 5), small biphasic spikes (cluster 2), and small monophasic spikes (cluster 4) were observed. To identify SSs in this cell accurately, further analysis was required (see Discussion). Small positive spikes sorted in cluster 1 could be ignored as artifacts.

**Fig. 5.**
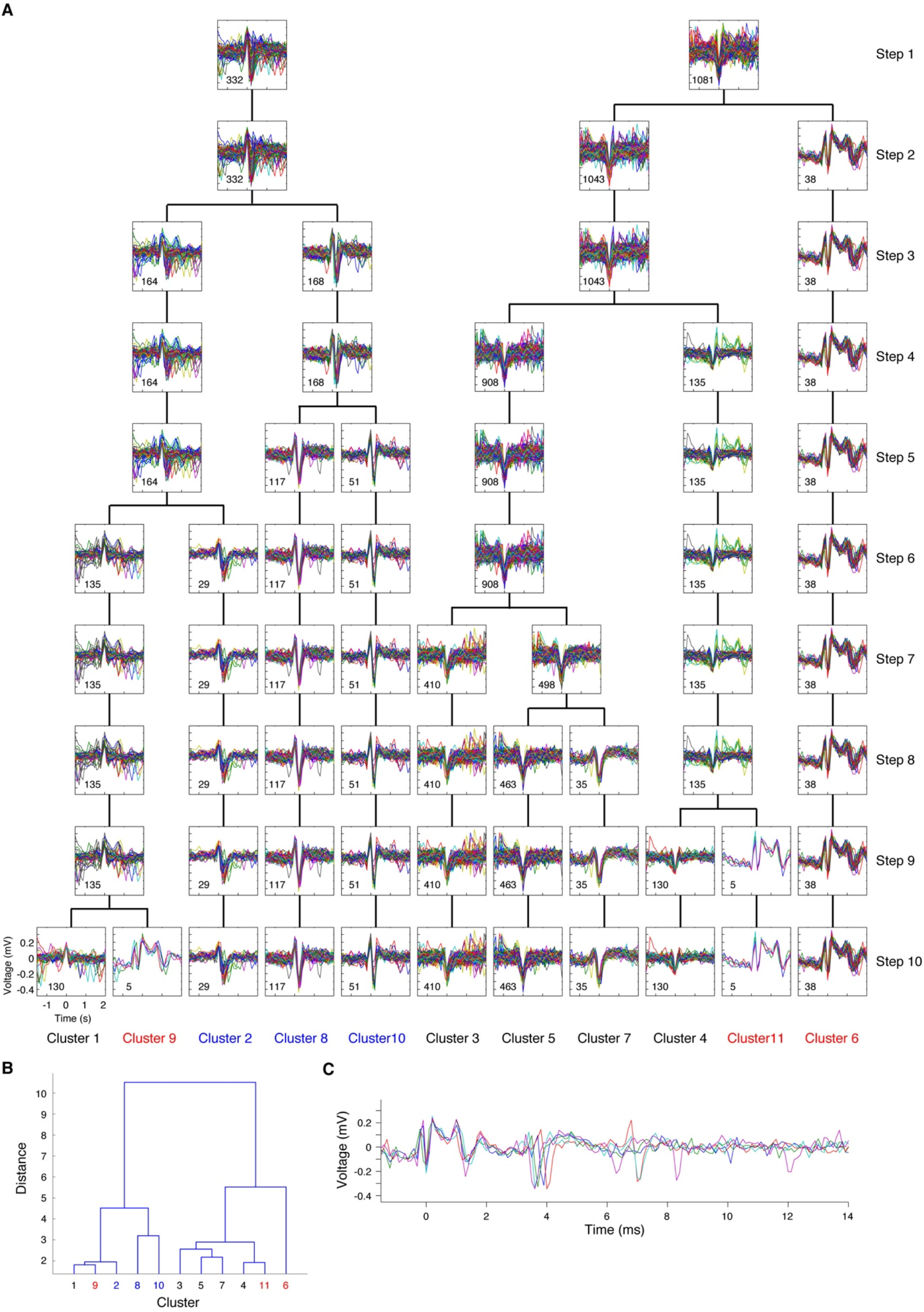
Spike sorting of data including CSs without initial negative peaks. (A) Results of hierarchical clustering that classified spikes detected in a 100-s recording into 11 clusters. The waveforms of individual spikes are superimposed with different colors in each plot. The number of spikes is shown in each panel. The clusters classified as CSs are shown in red (clusters 6, 9, and 11) and those classified as SSs are shown in blue (clusters 2, 8, and 10). (B) Dendrogram of hierarchical clustering. (C) The waveforms classified as CS are superimposed (n = 5).

### 3.4. Correction of the results of hierarchical clustering

After hierarchical clustering, the events categorized as SSs were evaluated by visual inspection (Fig. 2). Since the number of events categorized as SSs was too large to be inspected manually in many cases, a limited number of events were selected for visual inspection based on the following three criteria: (1) the variance of the waveform; (2) the temporal proximity to preceding events categorized as CS; and (3) the amplitude of the events. If CSs were misclassified as SSs, such occurrences would be selected using criterion (1), as their variances would be larger than the mean variance of the SS waveforms. If CS spikelets were misclassified as SSs, such events would be selected using criterion (2). If artifacts caused by misalignment of the event time were classified as SSs, such events would be selected using criterion (3). The selected events were evaluated visually and categorized into either SSs, CSs, or noise. After the correction, all events classified as CSs were evaluated by viewing their waveforms one-by-one (Fig. 2, see also supplemental materials).

### 3.5. Quantification of the performance of the spike-sorting algorithm

A representative result of simultaneous recordings of extracellular single-unit recordings and two-photon Ca^2+^ imaging of Purkinje cell dendrites from anesthetized mice is shown in Fig. 6. For this purpose, we used a fluorescence resonance energy transfer (FRET)-based ratiometric genetically-encoded Ca^2+^ indicator YC2.60 (Miyawaki et al., 1997; Nagai et al., 2004). During the recordings, FRET signals of YC2.60 repetitively exhibited a fast rise and slow decline. The occurrence of a CS labeled with a red circle was accompanied by a FRET signal increase (Fig. 6), indicating that the FRET signal captured the dendritic Ca^2+^ increase evoked by a single CS firing. Using the increase in FRET signal as an index of CS occurrences, we evaluated the spike-sorting algorithm based on hierarchical clustering as described above.

**Fig. 6.**
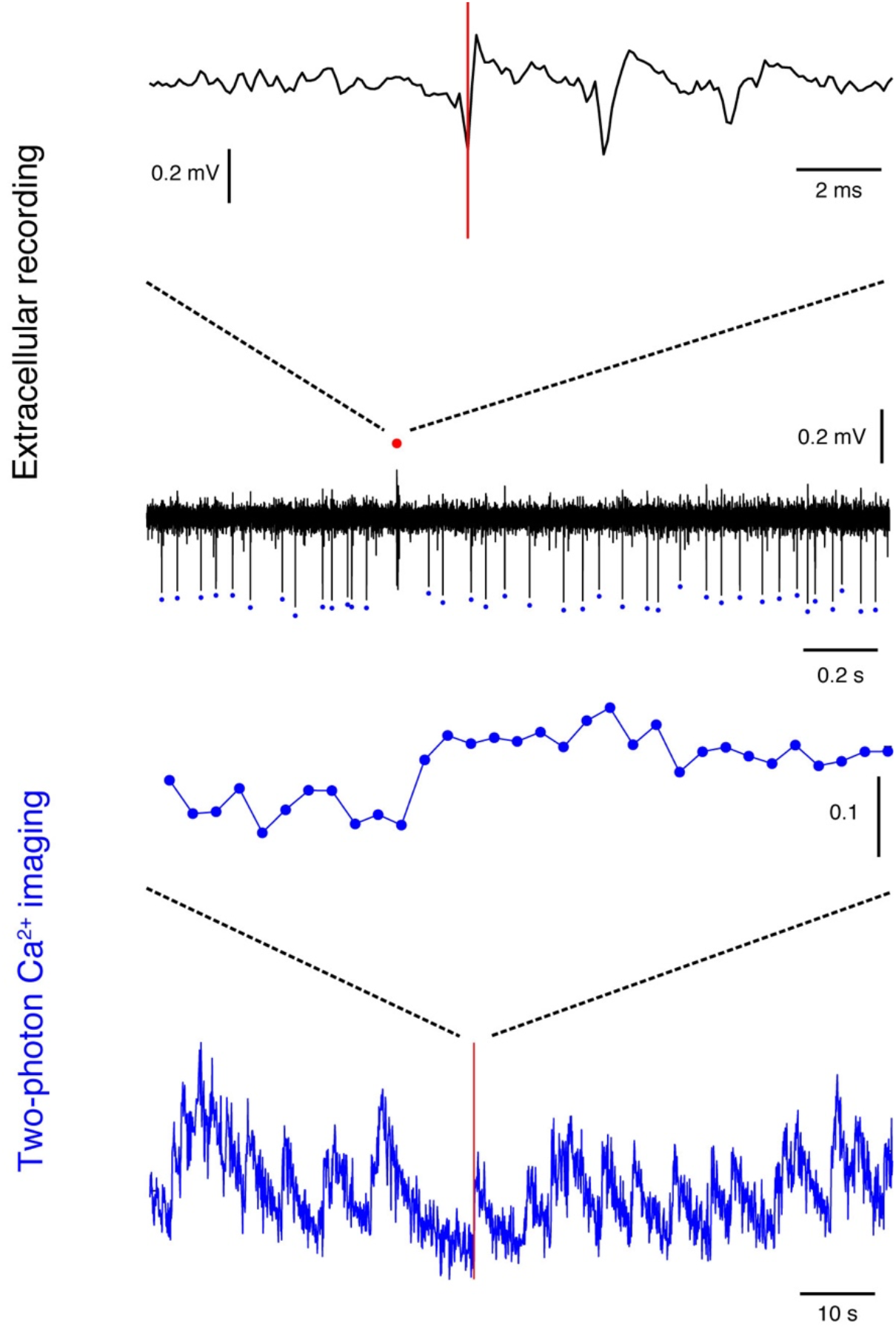
Comparison between extracellular single-unit recording and two-photon Ca^2+^ imaging data obtained simultaneously. The top panel shows a CS waveform on an extended time scale. The red vertical line indicates the onset time of CSs detected with *spike_hiclus.* The middle two panels show extracellular single-unit data (upper panel) and FRET signals of YC2.60 in Purkinje cell dendrites (lower panel) on the same time scale. The spike classified as a CS is shown in the red circle and those classified as SSs are shown in blue circles. The bottom panel shows all FRET signals obtained from the 100-s recording. The red vertical line indicates the spike onset time (same as the top panel). The sampling rate of the imaging data was 16 Hz.

The spike-sorting performance is summarized in Table 1. Among a total of 3,201 CSs detected with dendritic Ca^2+^ signals (the total recording time was 8,090 s from six cells in six mice), 2,826 (93.4%) were correctly sorted with hierarchical clustering. The number of misclassified spikes categorized as SSs was 154 (5.1%). Of these, 98 (3.2%) were selected during a manual correction of SS clusters and correctly categorized as CSs by visual inspection alone. Therefore, without simultaneously recorded two-photon Ca^2+^ imaging data, 96.6% of the CSs could be isolated by using the spike-sorting algorithm developed in this study. Examples of these initially misclassified and visually corrected spikes are shown in Fig. 7A and 7B. One reason for the misclassification is the misalignment of event time, as shown in Fig. 7A. Our algorithm estimates event time as the time that exhibits the minimal (or maximal) voltage signal within a range from –1.5 ms to +0.5 ms of the threshold-crossing time. This method worked well in most cases, but in a small number of events, there was an additional valley in the range, and thus, the event time was misaligned, as shown in Fig. 7A. These misaligned waveforms were not correctly categorized as CSs using hierarchical clustering. The other reason for the misclassification is the variability in the CS waveform itself. When the amplitude of the positive broad spike just following the initial negative potential was small, as shown in Fig. 7B, these spikes were sorted as SSs. These misclassified spikes shown in Fig. 7A and 7B were chosen automatically for visual inspection (the third stage in Fig. 2) because of their large variance. Once these spikes were inspected, they could easily be corrected as CSs by visual inspection.

**Fig. 7.**
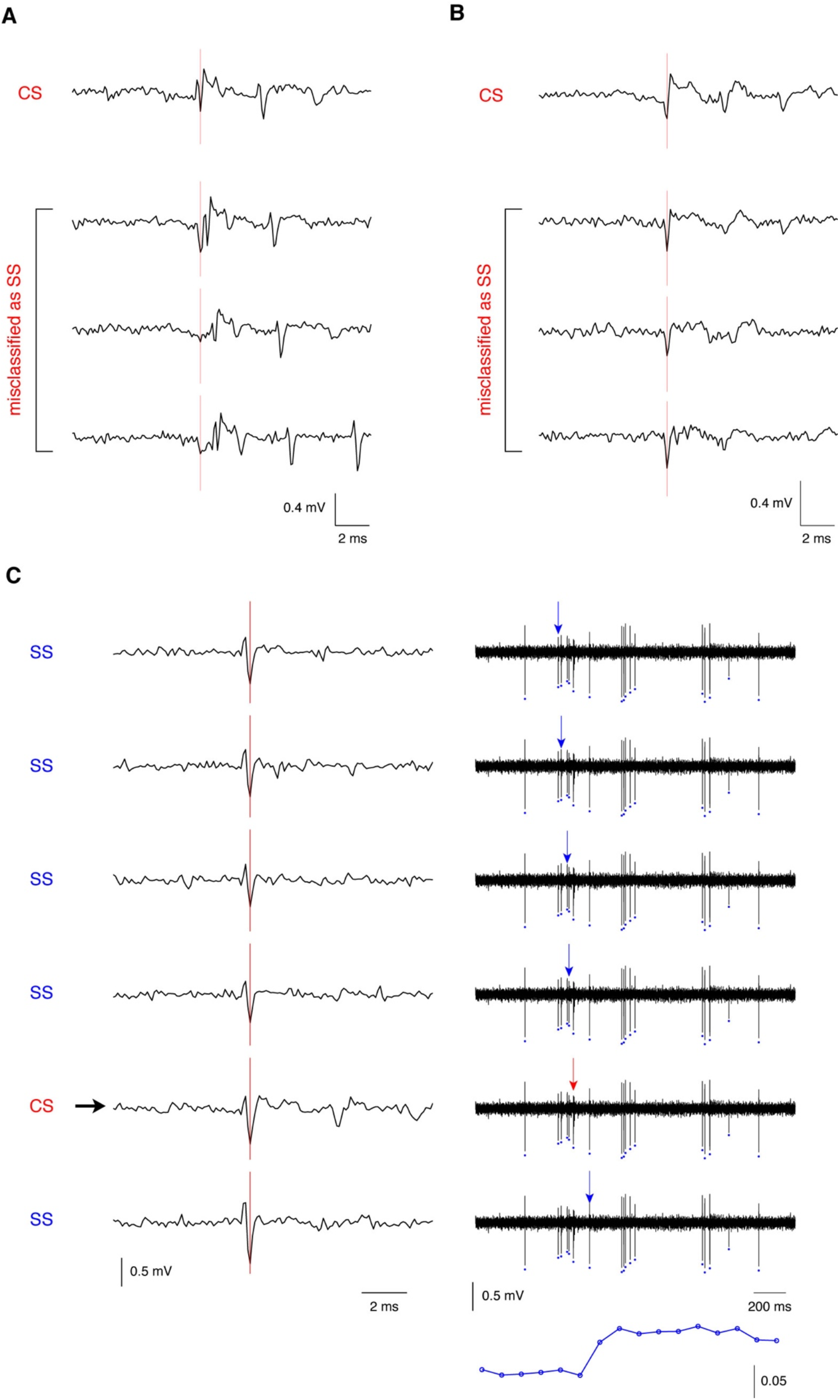
Misclassified CSs. (A) The top trace shows the waveform classified as a CS. The bottom three traces show waveforms of three different misclassified spikes caused by misalignments in the spike onset time (vertical red line). (B) The top trace shows the waveform classified as a CS. The bottom three traces show waveforms of three different misclassified spikes that exhibited smaller positive spikes. (C) The waveforms of six consecutive spikes, which were initially classified as SSs (left). The misclassified spike is marked in red. The estimated spike onset time is shown by the vertical red line. The temporal sequence of each spike is indicated by vertical arrows on the right-hand side of the plot. Simultaneously recorded FRET signals are shown at the bottom (blue). The black arrow on the left and the red arrow on the right indicate a misclassified CS.

**Table 1:** Summary of CS classification.

| Classification | Number of spikes | % |
| --- | --- | --- |
| 1. Correct classification |  |  |
| as CSs | 2,826 | 93.4 |
| 2. Misclassification |  |  |
| as SSs |  |  |
| - corrected by visual inspection | 98 | 3.2 |
| - corrected based on $\text{Ca}^{2+}$ imaging data | 56 | 1.9 |
| 3. Not detected | 42 | 1.4 |
| Total number of CSs | 3,021 | 100 |

Among the 154 misclassified spikes, 56 (1.9%) were not corrected by visual inspection of voltage signals alone, and a comparison with the two-photon Ca^2+^ imaging data was required. An example of these spikes is shown in Fig. 7C. FRET signals indicate the occurrence of a CS within the time range shown, in which all of the spikes were initially classified as SSs. A careful inspection of extracellular single-unit signals showed that a spike labeled with an arrow could correspond to a CS. Since the difference in the spike waveform was small among these spikes, identifying this CS should have been difficult without a comparison with the two-photon Ca^2+^ imaging data.

Among the total 3,021 CSs, 42 spikes (1.4%) were marked as neither CSs nor SSs because the amplitude was smaller than the threshold, or because small spikes sorted by hierarchical clustering, i.e., cluster 2 in Fig. 3A and cluster 4 in Fig. 5A, were ignored as noise. A example of such missed spikes is shown in Fig. 8. FRET signals indicate the occurrence of CSs in both cases, but typical CS waveforms were not found in the ranges shown. Small voltage signals around 20.6 s (Fig. 8A) and 60.9 s (Fig. 8B) may have corresponded to CSs in the recording range shown. These results suggest that CS waveforms exhibit large variations, that around 2% of CSs are misclassified as SSs even after manual correction, and that around 1% of CSs exhibit a small nonstandard waveform that is undetectable using the spike-sorting algorithm.

**Fig. 8.**
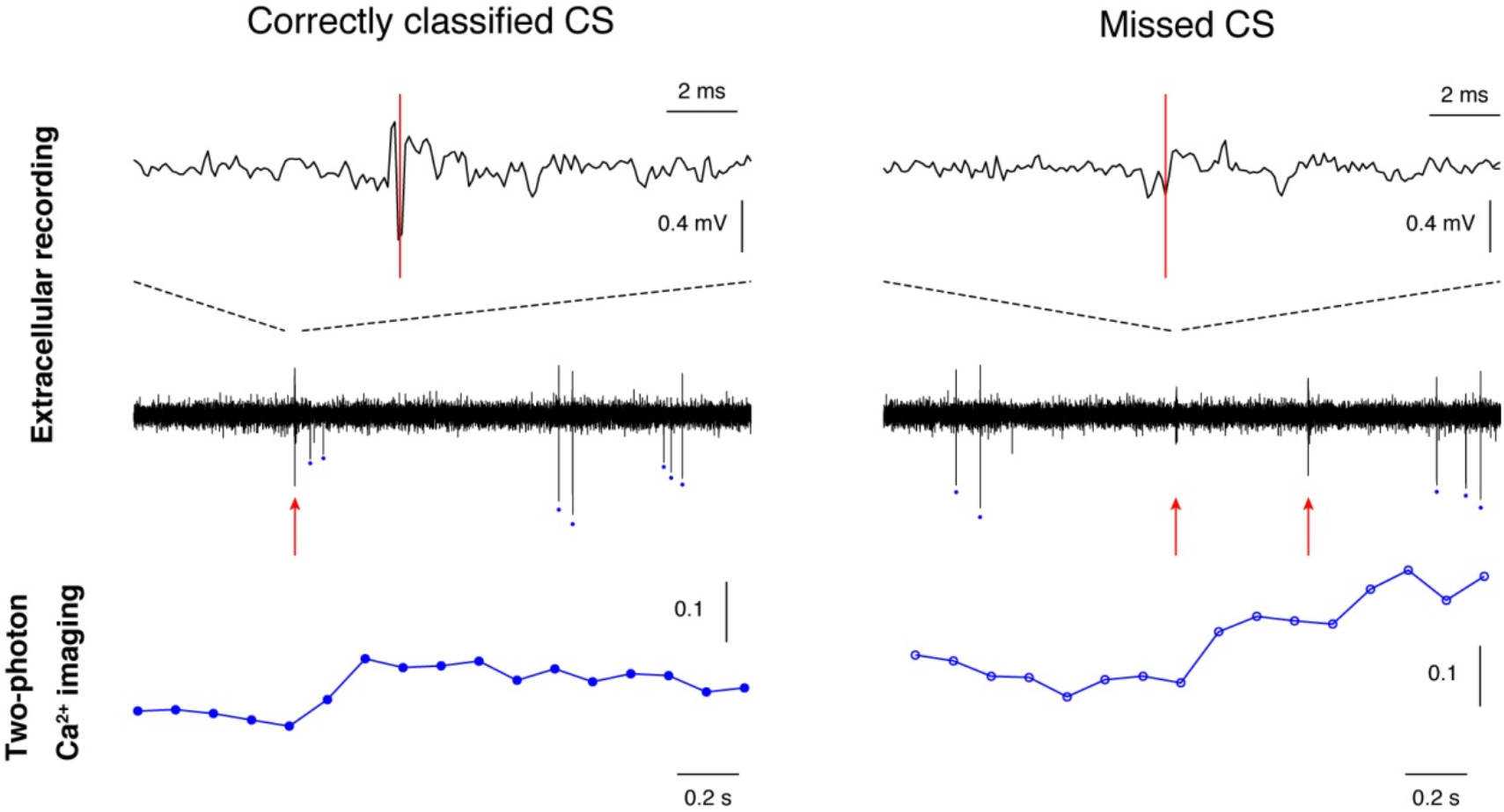
Missed CSs. The first row shows the waveforms of a spike classified as a CS (left) and a missed spike (right) in the same data set. The second row shows extracellular single-unit data around the aforementioned spikes. CSs are indicated with red arrows and SSs are indicated with blue circles. The bottom row shows concurrently recorded FRET signals of Purkinje cell dendrites. The sampling rate of the imaging data was 8 Hz.

### 3.6. Inter-spike interval of CSs

An inter-spike interval histogram of the detected CSs (total of 3,021 spikes) from fentanyl-anesthetized mice is shown in Fig. 9. The minimal interval was 0.0106 s, corresponding to 94.3 Hz.

**Fig. 9.**
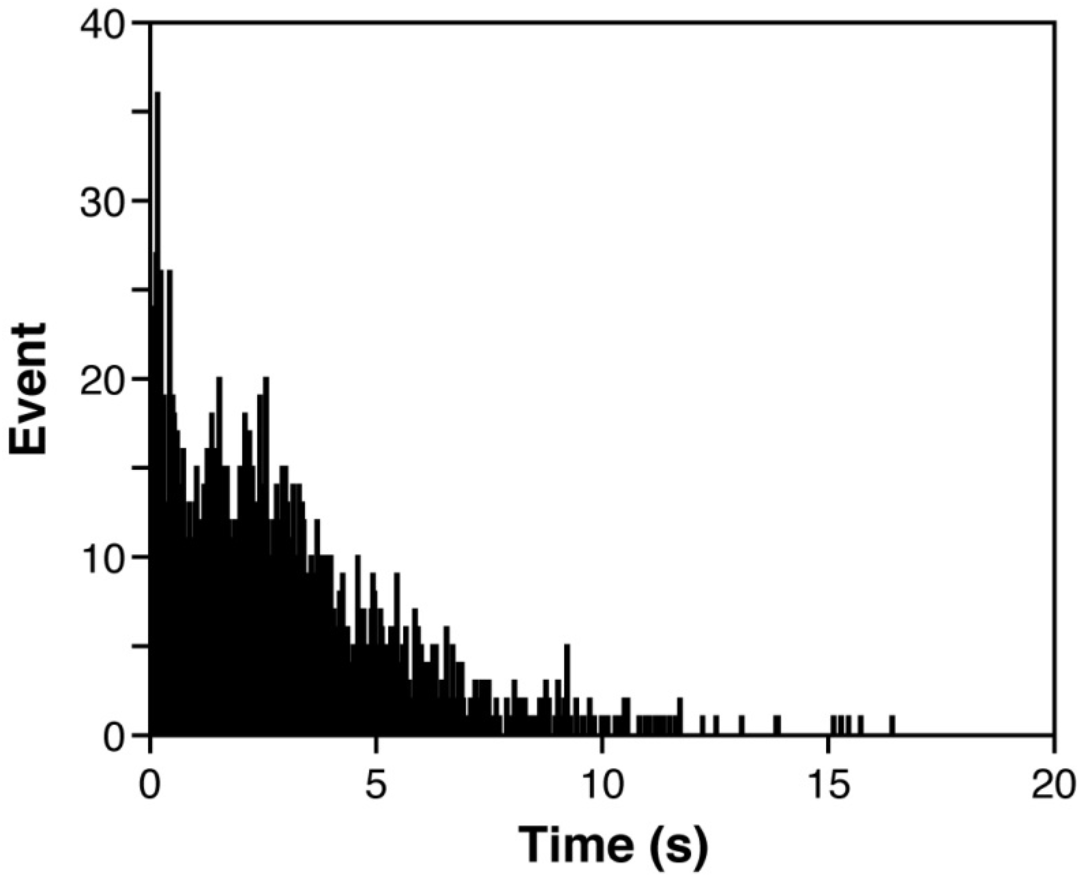
Inter-spike interval histogram of CSs. Inter-spike interval histogram of CSs identified from extracellular single-unit recordings (total of 3,021 spikes) from fentanyl-anesthetized mice using the spike-sorting algorithm.

## 4. Discussion

In this study, we developed a spike-sorting method based on divisive hierarchical clustering to categorize two types of spikes (SSs and CSs) from extracellular single-unit recording data obtained from cerebellar Purkinje cells. Simultaneous *in vivo* dendritic Ca^2+^ imaging was used for CS detection to evaluate the performance of the spike-sorting method. Previous *in vitro* measurements have shown that artificially strong activations of parallel fibers can evoke localized Ca^2+^ rises in Purkinje cell dendrites in acute cerebellar slices (Hartell, 1996). However, massive increases in Ca^2+^ across dendrites should rarely be elicited with parallel fiber inputs under physiological conditions. We used the increase in the FRET signal as an index of the occurrence of CSs in living mice and demonstrated that a spike-sorting algorithm based on hierarchical clustering could identify 96.6% of CSs from extracellular single-unit recordings.

A quantification of the accuracy of spike sorting based on ground-truth data was carried out by simultaneous recordings of both intracellular and extracellular signals from the same hippocampal neurons (Harris et al., 2000). The authors showed that when the first three principal components of extracellular spike waveforms are used to construct the feature vectors, the main cause of the separation error resulting in false negative and false positive spikes ranging from 0%to 30% is the inability of human operators to mark boundaries accurately in the high-dimensional feature space. The authors also showed that a semiautomatic classification system in which boundaries are determined automatically and the operator merges or splits clusters manually can reduce errors in the 0%–8% range. Our algorithm uses the extracellular spike waveforms themselves as feature vectors and automatically performs spike classification with hierarchical clustering. The operator splits the clusters by manually increasing the number of hierarchical clusters and finally selects those corresponding to SSs and CSs. Comparisons between the extracellular single-unit and two-photon Ca^2+^ imaging data showed that the main cause of detection errors (false negatives) of CSs with the algorithm was intrinsic fluctuations of the CS waveform, rather than operation errors. We found that spontaneous Purkinje cell firing in the anaesthetized mouse cerebellum involves around 2% of CSs whose waveforms are indistinguishable to those of SSs with the algorithm used, and around 1% of CSs whose waveforms are undetectable under the conditions used.

### 4.1. Variability in the CS waveforms

A climbing fiber excites cerebellar Purkinje cells through hundreds of widespread dendritic glutamatergic synapses, triggering CSs (Eccles et al., 1966) and dendritic Ca^2+^ spikes that can propagate toward the soma (Fujita, 1968; Llinás and Sugimori, 1980). The first spike, as well as all spikelets within the CS, originates from the axon of the Purkinje cell, and the dendritic Ca^2+^ spikes play a minor role in generating the CSs (Davie et al., 2008). Two-photon Ca^2+^ imaging measures widespread dendritic Ca^2+^ increases (Miyakawa et al., 1992; Ross and Werman, 1987) mediated by P/Q-type (Llinás et al., 1989) and T-type (Watanabe et al., 1998) voltage-gated Ca^2+^ channels activated by climbing fiber input. The extracellular electrode captures extracellular voltage changes resulting from the spiking activity of nearby neurons (Gold et al., 2006). In this study, the tip of the electrode was strictly targeted to the Purkinje soma, which closes the CS initiation site, but stereotypic CS waveforms were not detected in around 3% of the dendritic Ca^2+^ increases detected using two-photon Ca^2+^ imaging. Therefore, it remains unknown whether the CS generation in a cell or the extracellular action potential detection failed in these cases. Kitamura and Häusser (2011) showed that 95.0% ± 2.1% of spontaneous dendritic Ca^2+^ signals were associated with CSs in rat cerebellar Purkinje cells. Because they employed the *in vivo* somatic and dendritic patch-clamp technique to track the firing of Purkinje cells, widespread dendritic Ca^2+^ increases occurred without the chance to generate CSs in the axon to a smaller extent (approximately a few percent of the total events). We cannot exclude the possibility of spontaneous dendritic Ca^2+^ rises without climbing fiber inputs in our measurements, but if climbing fiber inputs occasionally fail to induce stereotypic CS waveforms, even when dendritic Ca^2+^ rises are elicited, *in vivo* extracellular recordings slightly underestimate the occurrence of climbing fiber inputs.

### 4.2. Identification of SSs

It is difficult to identify SSs when there are multiple waveforms that are not categorized as CSs, as shown in Fig. 5. One explanation for the variability in waveforms is that extracellular recordings contain signals originating from multiple cells. In the present measurements, as mentioned above, the tip of the electrode was strictly targeted to the Purkinje soma within the Purkinje cell layer under visual inspection. That is, neighboring Purkinje cells are a major candidate for the source of multiple waveforms recorded, rather than cells in other layers, such as granule cells in the granule cell layer and interneurons in the molecular layer. Considering that nearly 100% of the CS waveforms were matched with concurrently measured dendritic Ca^2+^ increases, contaminated signals should have been SSs generated by neighboring Purkinje cells that exhibit no CSs. No CS Purkinje cells were previously detected under anesthesia with *in vivo* two-photon Ca^2+^ imaging (Mukamel et al., 2009). Another possibility is that a single Purkinje cell exhibits multiple SS waveforms. Variations in spike amplitudes produced by cardiovascular movements of the cerebellum have been detected in extracellular signals (Brookhart et al., 1950).

The effects of the interactions between SSs and CSs have been reported (Bell and Grimm, 1969; Granit and Phillips, 1956; Thach, 1967). All examined Purkinje cells show a pause in the SS train lasting from 10 to >100 ms after the occurrence of a CS (Sato et al., 1992). After a pause, some cells exhibit transient facilitation or a reduction in the SS firing frequency (Sato et al., 1992). Therefore, if the firing pattern of spikes decreases after CS firing, such spikes must be SSs originating from the same Purkinje cell. Examples of such an analysis are shown in Fig. S7. A raster plot (upper panel) and a peri-CS time histogram (bottom panel) of the biphasic spike (Fig. S7A), which was originally classified into clusters 8 and 10 of the data presented in Fig. 5, are shown in Fig. S7A and S7B. This spike clearly exhibited a pause after CSs, indicating that it corresponds to SSs. By contrast, the firing rates of the monophasic spikes (Fig. S7C), which were originally classified into clusters 3 and 5 (Fig. 5), were independent of the occurrence of CSs (Fig. S7D). It should be noted that we cannot evaluate whether the spikes classified into clusters 3 and 5 were SSs in this case because some cells only exhibited a brief pause (approximately 20 ms) in the SS train after the CS without any changes in SS firing rates after the pause (Sato et al., 1992). In the spike-sorting software developed in this study, any spikes occurring within 15 ms after the onset of CSs are usually ignored to avoid false spikelet detection within CS waveforms as independent spikes. Therefore, the existence of a pause alone cannot be used to identify SSs in these cells, and a different criterion is required. In practice, the shapes of SSs and CSs originating from the same Purkinje cell simultaneously change depending on the position of the recording electrode. This information could be expected to help identify SSs for off-line analysis.

## 5. Conclusion

For extracellular records from cerebellar Purkinje cells, we developed a spike-sorting method based on hierarchical clustering. The quantitative evaluation of the spike-sorting algorithm with the use of simultaneously recorded two-photon Ca^2+^ imaging data showed that it can identify 96.6% of CSs from extracellular single-unit recordings. Because this method is semiautomatic, operator supervision is required for several processes, such as (1) determining the number and polarity of threshold(s) to detect events, (2) determining the threshold value(s), (3) determining the number of hierarchical clusters, (4) assigning SSs and CSs to each cluster, (5) correcting events assigned as SSs or CSs, and (6) determining the number of clusters for sub-classification of CSs. In the future, an efficient application of the deep learning methodology based on ground-truth data obtained by using the spike-sorting method could be expected to help develop a fully automatic, unsupervised spike-sorting algorithm for extracellular recordings from cerebellar Purkinje cells.

## Acknowledgments

We wish to thank Dr. Yoshikazu Isomura, Dr. Takahiro Ishikawa, and Dr. Michael London for technical help with the extracellular single-unit recordings. This work was supported by a Japan Ministry of Education, Culture, Sports, Science and Technology Grant-in-Aid for Scientific Research (C) (24500476), JST CREST (JPMJCR1851 and JPMJCR1921).

