## Supplemental materials for "A spike-sorting method based on hierarchical clustering to discriminate extracellularly recorded simple spikes and complex spikes from cerebellar Purkinje cells"

**Supplementary materials**

***spike_hiclus*, spike-sorting software using hierarchical clustering**

For the off-line spike sorting of extracellular single-unit data recorded from cerebellar Purkinje cells, the spike-sorting software *spike_hiclus* which is composed of four class files, was implemented in MATLAB. The software has four steps to isolate SSs and CSs: (1) filtering, (2) spike detection, (3) spike sorting, and (4) spike classification correction.

1. Filtering

The extracellular single-unit data are first filtered with a noncausal bandpass filter. The default settings for the lower and upper cutoff frequencies are 300 and 3,000 Hz, respectively. The MATLAB command of '*filtfilt*' in the Signal Processing Toolbox can be used as a noncausal filter.

2. Spike detection

To refine the spike-detection method, a continuous 200-ms time window with the highest variance is automatically selected (Fig. S1A). Spikes can be detected with a positive threshold, a negative threshold, or both depending on the spike waveforms (Fig. S1B). Moreover, in this step, the spike waveform alignment method is chosen (Fig. S1B). For spikes detected with a positive threshold or both positive and negative thresholds, waveforms can be aligned to the time of either the maximal or minimal voltage signal within –1.5 ms to +0.5 ms relative to the time of crossing the threshold. For spikes detected with a negative threshold, waveforms are always aligned to the time of the minimal voltage signal within –1.5 ms to +0.5 ms relative to the time of crossing the threshold.

Peak amplitude is determined with the MATLAB command '*findpeaks*' provided in Signal Processing Toolbox. Because only positive peaks can be detected with *findpeaks*, the voltage signal signs are changed when a negative threshold is applied. Among the following three choices, a threshold value for spike detection is chosen: (1) the half difference between the centroids of two clusters of peak amplitudes determined using k-means clustering, (2) the half amplitude of the largest (or 21st or 41st) peak value, or (3) the fivefold standard deviation of the baselines, calculated with Equation (1) described in the main text. Examples of threshold determination for biphasic positive–negative potentials with a large positive peak, biphasic potentials with a small positive peak, and a monophasic negative peak are shown in Figs. S2–S4, respectively. In these figures, the results of k-means clustering of the peak heights are shown in A and B. The peak height of the top 100 largest peaks is shown in C. The waveforms of the 1st- (blue), 21st- (red), and 41st- (green) largest peaks are shown in D. Overall event counts and firing rates using different thresholds are shown in E and F, respectively. Five threshold values are superimposed on the histogram of peak heights in G. The user interface for choosing the threshold is shown in H. The estimation of half of the amplitude of spikes is difficult if artifacts with large amplitudes are involved in the data. Therefore, to exclude artifacts, the waveforms of the 1st- (blue), 21st- (red), and 41st- (green) largest peaks are shown in D.

When spikes show a large positive transient with little baseline noise, k-means clustering performs well (Fig. S2A,B), and the choice of (1) is preferred (Fig. S2G). However, in many cases, k-means clustering cannot isolate spikes from baseline noise (Figs. S3A,B and S4A,B), and thus the choice of (2) or (3) is appropriate. When multiple spike shapes are involved in the record (see below), the peak amplitude exhibits a broad distribution, and a threshold determined based on the largest peak amplitude does not work well to detect spikes. The choice of (3) is appropriate for these records. However, since small noises can be excluded during the subsequent hierarchical clustering (see below), choosing a lower threshold value is generally preferable to reducing the number of missed spikes.

After the determination of threshold crossing times, the event time is defined as follows. For events detected with a positive threshold, the event time is defined as the time of maximal or minimal voltage signals within –1.5 ms to +0.5 ms of the time of threshold crossing depending on the alignment method. For events detected with a negative threshold, the event time is defined as the time of minimal voltage signals. The events are merged if both negative and positive thresholds are used. To avoid multiple selections of single events in the merged data, only the events that were apart by an interval (default: 1.5 ms) from the preceding event are adapted. Then, waveforms are constructed based on the event time by taking from –1.5 ms to +2 ms of the filtered voltage signals for each event for application to the hierarchical clustering.

3. Spike sorting

Hierarchical clustering is performed using MATLAB with the commands ‘*pdist*’, ‘*linkage*’, and ‘*dendrogram*’ provided in the Statistics and Machine Learning Toolbox. The process was started with two clusters, and the number of clusters can be increased to the maximum number determined (default setting: 11) in an interactive manner. Also, the number of clusters can be reduced interactively. Thus, by viewing all the spikes superimposed in each cluster, an optimal cluster number can be determined by changing the cluster number back and forth. Once the CS clusters are selected, the hierarchical clustering can be terminated. It is also possible to select clusters to be discarded.

4. Correction of spike classification

In many cases, the number of spikes classified as SSs is too large to be inspected manually. Therefore, we select spikes for visual inspection based on the following three criteria: (1) waveform variance; (2) temporal proximity to the previous spike, which is classified as a CS; and (3) spike amplitude. Spikes with a large variance are selected as follows. The average waveform of spikes temporally categorized as SSs is computed. The difference between the average waveform and all the spikes temporally categorized as CSs is computed. The minimum value among these differences is used as the threshold. Then, the difference between the average waveform and all the spikes temporally categorized as SSs is computed, and the spikes with a difference larger than the threshold are selected.

The software automatically selects spikes with waveform difference larger than the threshold, events close to a previous spike categorized as a CS (< 16 ms), and spikes with an amplitude smaller than the event detection threshold. The selected spikes are displayed with two different time scales, as shown in Fig. S5. The operator assigns them as either SSs, CSs, or noise based on visual inspection. For example, in Fig. S5, the spike is selected with its large variance caused by the closeness of the subsequent CS and should be assigned as an SS. After the correction of SSs, the correction of spikes temporally categorized as CSs is performed. In this step, all spikes are applied for visual inspection to be assigned as either CSs, SSs, or noise. After SSs and CSs are confirmed, the spike times of both are stored into text files.

After the spike classification is corrected, spike time alignment is performed on both SSs and CSs, since misaligned spikes (i.e., Fig. 7A) may be involved in either SS or CS clusters. SSs are aligned to their minimum. For the alignment of CSs, a template waveform is selected.

5. Sub-classification of CSs

CSs can be further classified depending on the shape of spikelets by using the same hierarchical clustering algorithm (Fig. S6). The number of subclasses can be increased in an interactive manner (Fig. S6A). Color-coded CS waveforms are displayed to evaluate the appropriateness of the classification (Fig. 6A). A dendrogram is also shown to confirm the similarity between subclasses (Fig. S6B).

### Supplemental Figures

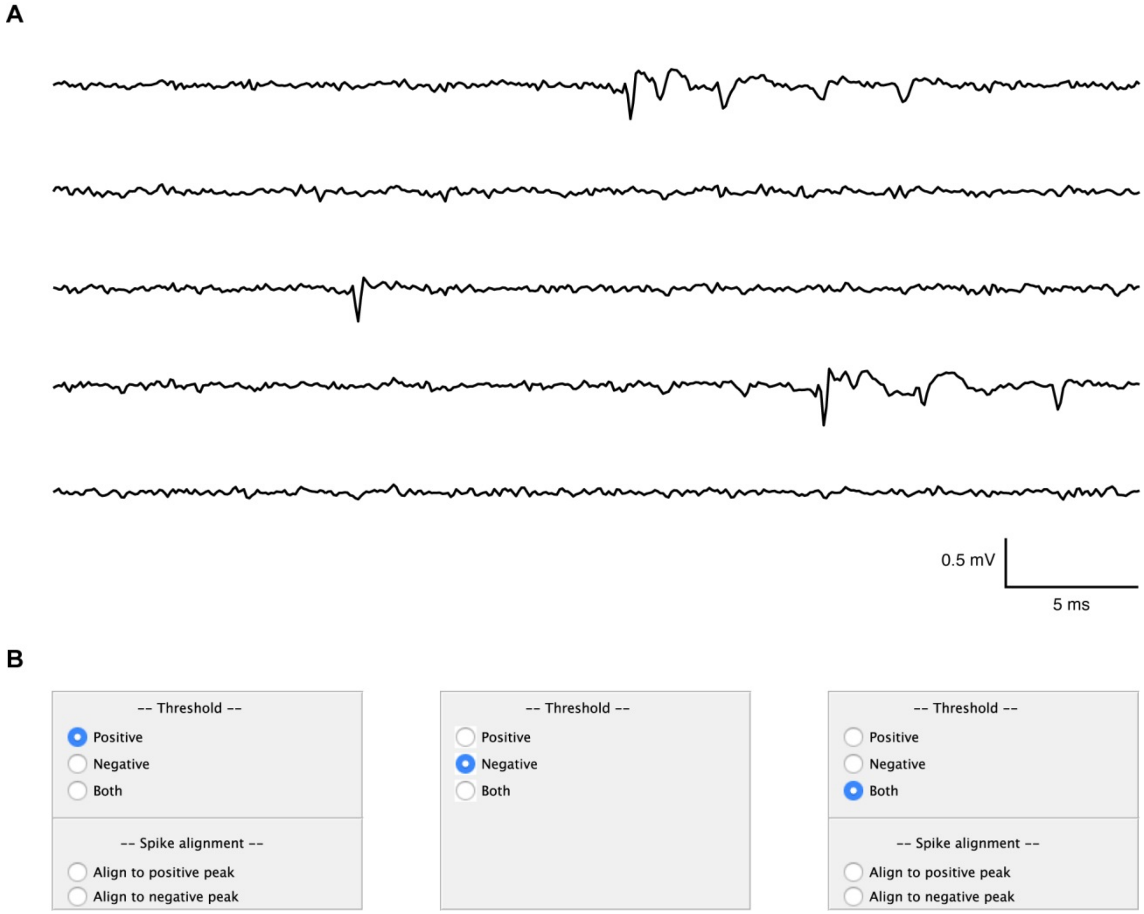


**Fig. S1. Choice of spike detection method and alignment polarity of spikes.** (A) An sample of 200 ms of contiguous records showing the highest variance among the entire filtered extracellular single-unit record. (B) Graphical user interface panels are used to select threshold(s) for detecting spikes and the alignment polarity of spikes detected.

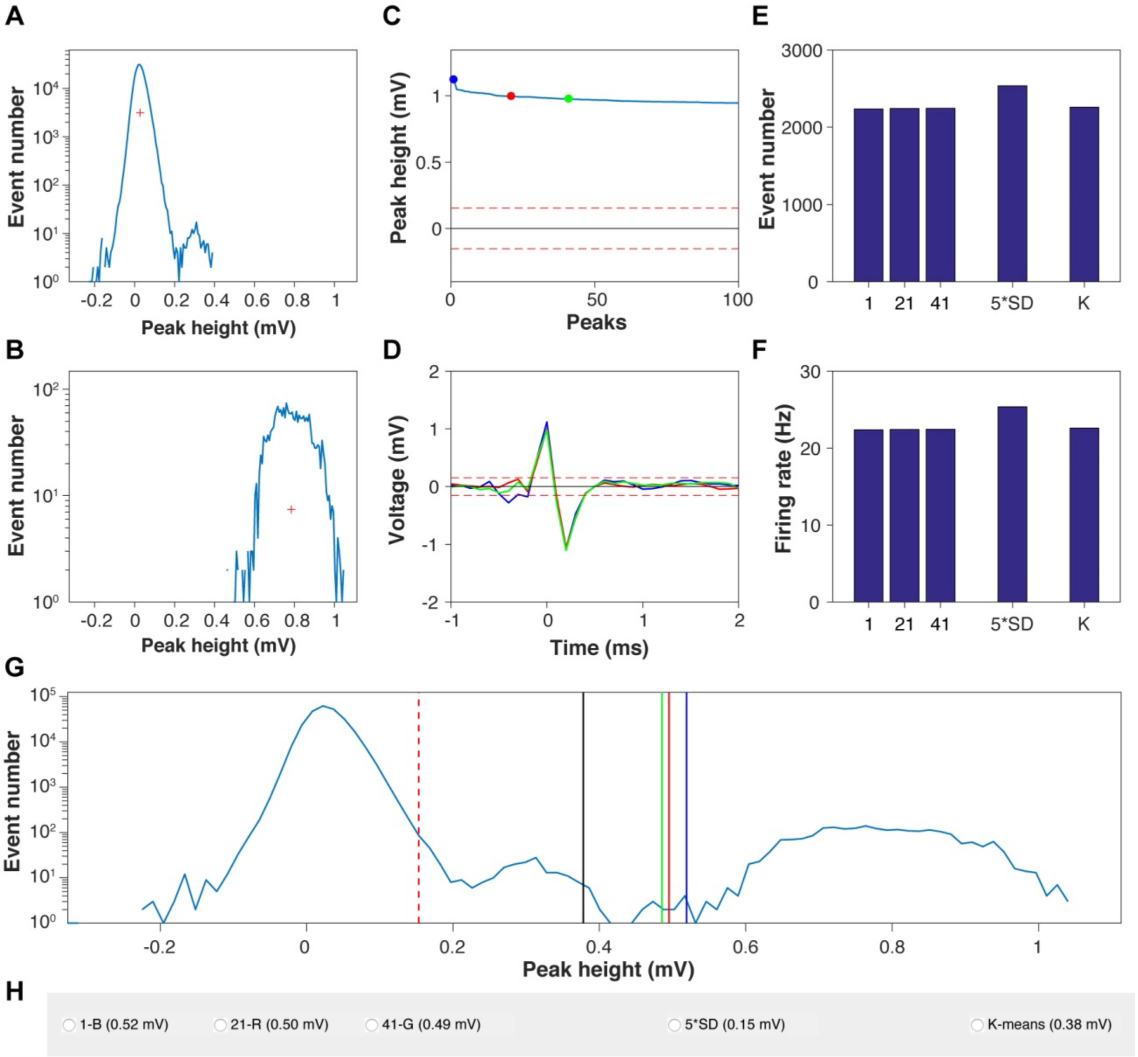


**Fig. S2. Selecting the spike detection threshold for biphasic spikes.** (A and B) Histograms of the peak amplitudes of two clusters divided by k-means clustering. Red crosses indicate the centroid of each cluster. (C) Peak heights of the top 100 spikes sorted in descending order. The 1st, 21st, and 41st spikes are indicated with blue, red, and green circles, respectively. The red dashed lines indicate the fivefold standard deviation of the baseline. (D) Spike waveforms of the 1st (blue), 21st (red), and 41st (green) spikes. The red dashed lines indicate the fivefold standard deviation of the baseline. (E and F) Numbers of spikes (E) and firing rates (F) using a different threshold. 1: half amplitude of the 1st spike; 21: half amplitude of the 21st spike; 41: half amplitude of the 41st spike; 5*SD: fivefold standard deviation of the baseline; K: k-means clustering. (G) Each threshold is superimposed on the distribution of the peak amplitude. Blue: half amplitude of the 1st spike; red: half amplitude of the 21st spike; green: half amplitude of the 41st spike; red dashed line: fivefold standard deviation of the baseline; black: k-means clustering. (I) Graphical user interface panel for choosing a threshold. Threshold values are indicated in parentheses. These data are shown in Fig. 4.

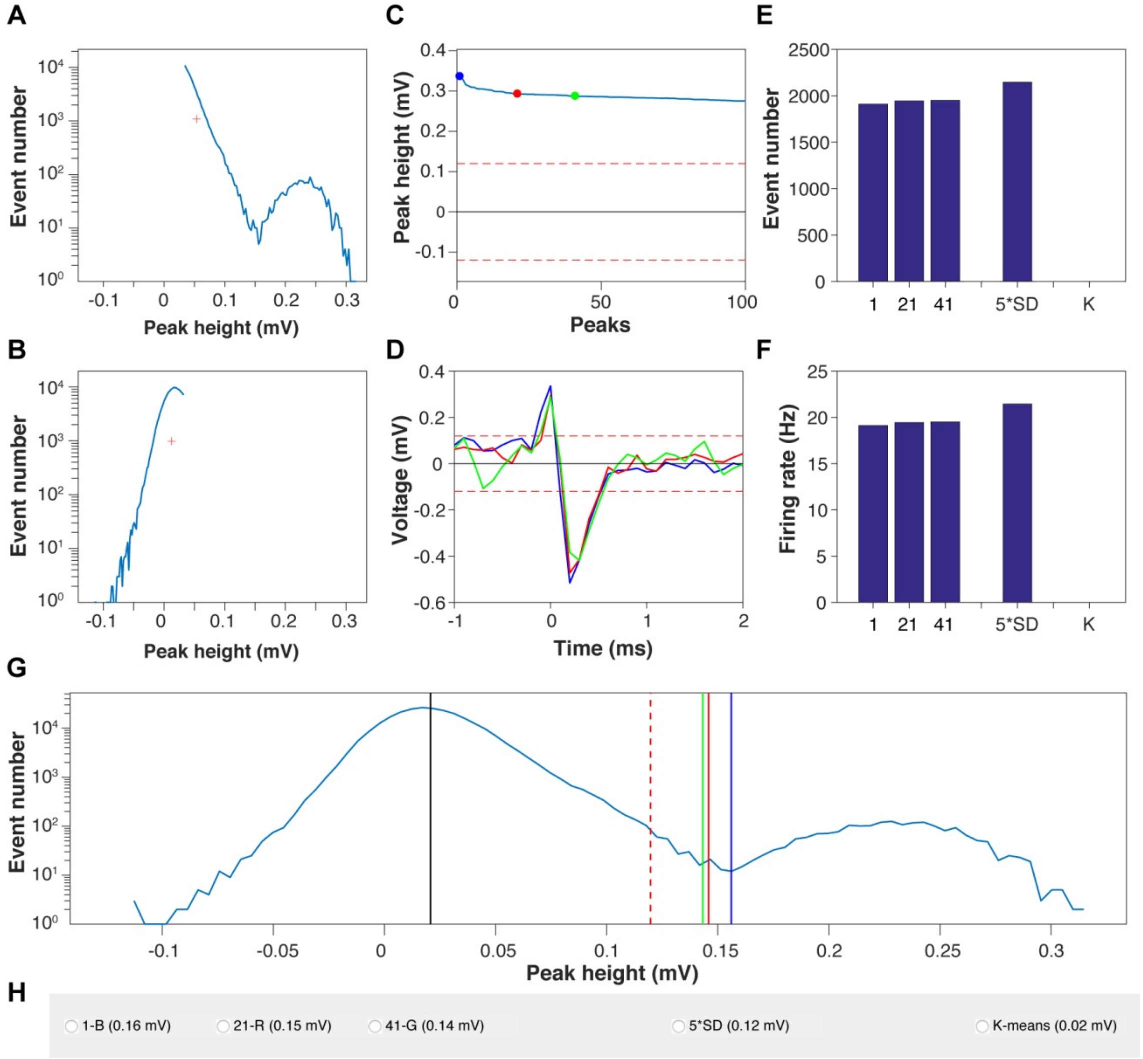


**Fig. S3. Selecting the spike detection threshold for biphasic spikes with a small positive peak.** (A and B) Histograms of the peak amplitudes of two clusters divided by k-means clustering. Red crosses indicate the centroid of each cluster. (C) Peak heights of the top 100 spikes sorted in descending order. The 1st, 21st, and 41st spikes are indicated with blue, red, and green circles, respectively. The red dashed lines indicate the fivefold standard deviation of the baseline. (D) Spike waveforms of the 1st (blue), 21st (red), and 41st (green) spikes. The red dashed lines indicate the fivefold standard deviation of the baseline. (E and F) Numbers of spikes (E) and firing rates (F) using a different threshold. 1: half amplitude of the 1st spike; 21: half amplitude of the 21st spike; 41: half amplitude of the 41st spike; 5*SD: fivefold standard deviation of the baseline; K: k-means clustering. (G) Each threshold is superimposed on the distribution of the peak amplitude. Blue: half amplitude of the 1st spike; red: half amplitude of the 21st spike; green: half amplitude of the 41st spike; red dashed line: fivefold standard deviation of the baseline; black: k-means clustering. (I) Graphical user interface panel for choosing a threshold. Threshold values are indicated in parentheses. These data are shown in Fig. 5.

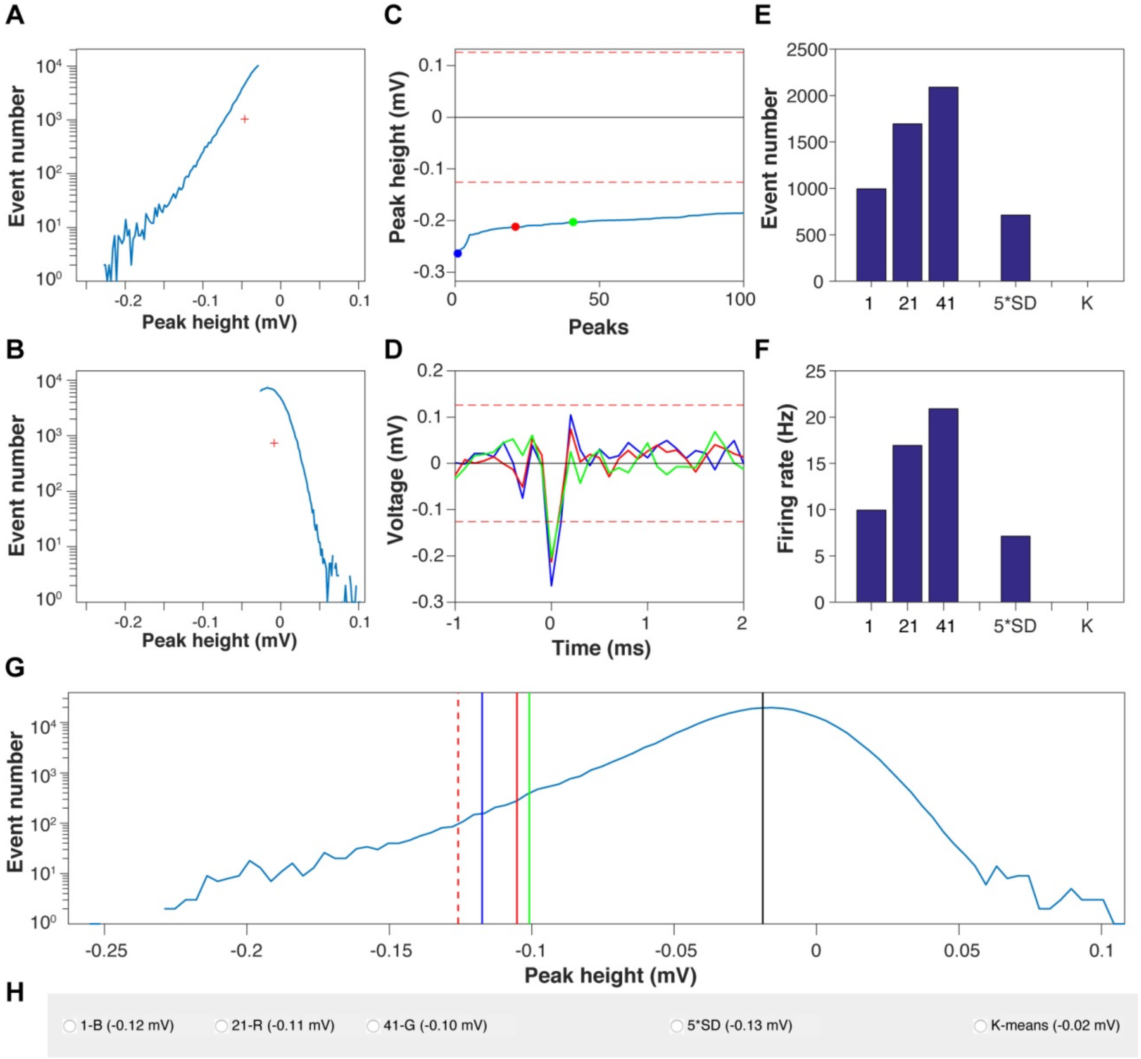

**Fig. S4. Selecting the spike detection threshold for monophasic negative peaks.** (A and B) Histograms of the peak amplitudes of two clusters divided by k-means clustering. Red crosses indicate the centroid of each cluster. (C) Peak heights of the top 100 spikes sorted in descending order. The 1st, 21st, and 41st spikes are indicated with blue, red, and green circles, respectively. The red dashed lines indicate the fivefold standard deviation of the baseline. (D) Spike waveforms of the 1st (blue), 21st (red), and 41st (green) spikes. The red dashed lines indicate the fivefold standard deviation of the baseline. (E and F) Numbers of spikes (E) and firing rates (F) using a different threshold. 1: half amplitude of the 1st spike; 21: half amplitude of the 21st spike; 41: half amplitude of the 41st spike; 5*SD: fivefold standard deviation of the baseline; K: k-means clustering. (G) Each threshold is superimposed on the distribution of the peak amplitude. Blue: half amplitude of the 1st spike; red: half amplitude of the 21st spike; green: half amplitude of the 41st spike; red dashed line: fivefold standard deviation of the baseline; black: k-means clustering. (I) Graphical user interface panel for choosing a threshold. Threshold values are indicated in parentheses. These data are shown in Fig. 3.

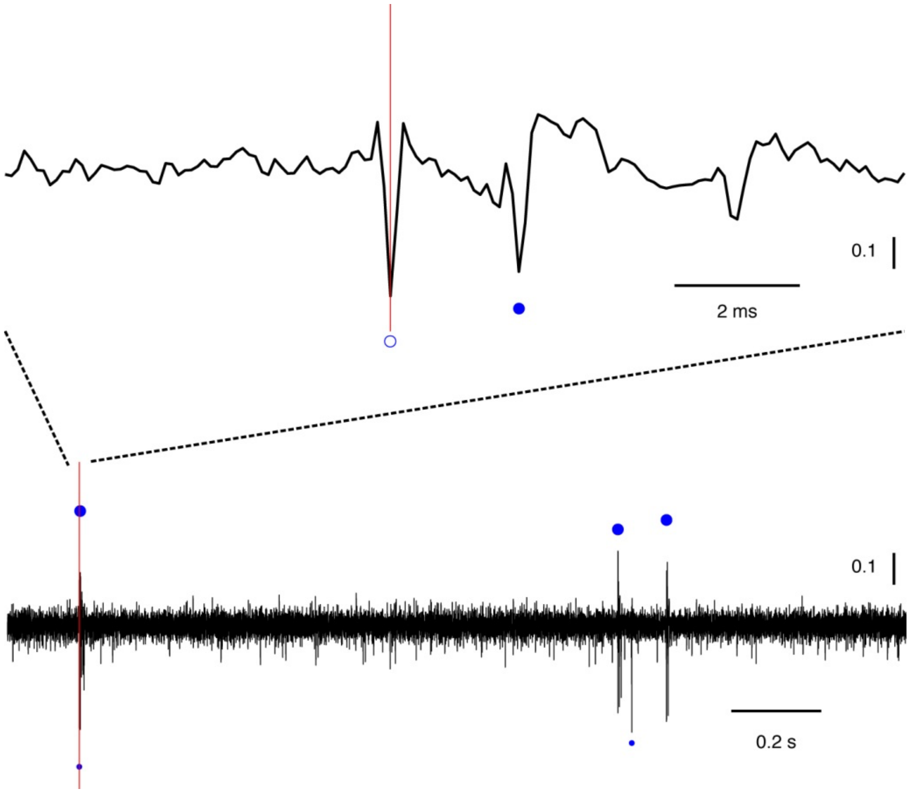

**Fig. S5. Example of the evaluation of a spike classified as an SS.** In the evaluation, a spike (vertical red line) is shown at two different time scales. A spike classified as an SS is indicated with an open circle, and a spike classified as a CS is indicated with a filled circle (top). Spikes classified as an SS are indicated with small circles below the trace, and spikes classified as a CS are indicated with large circles above the trace (bottom).

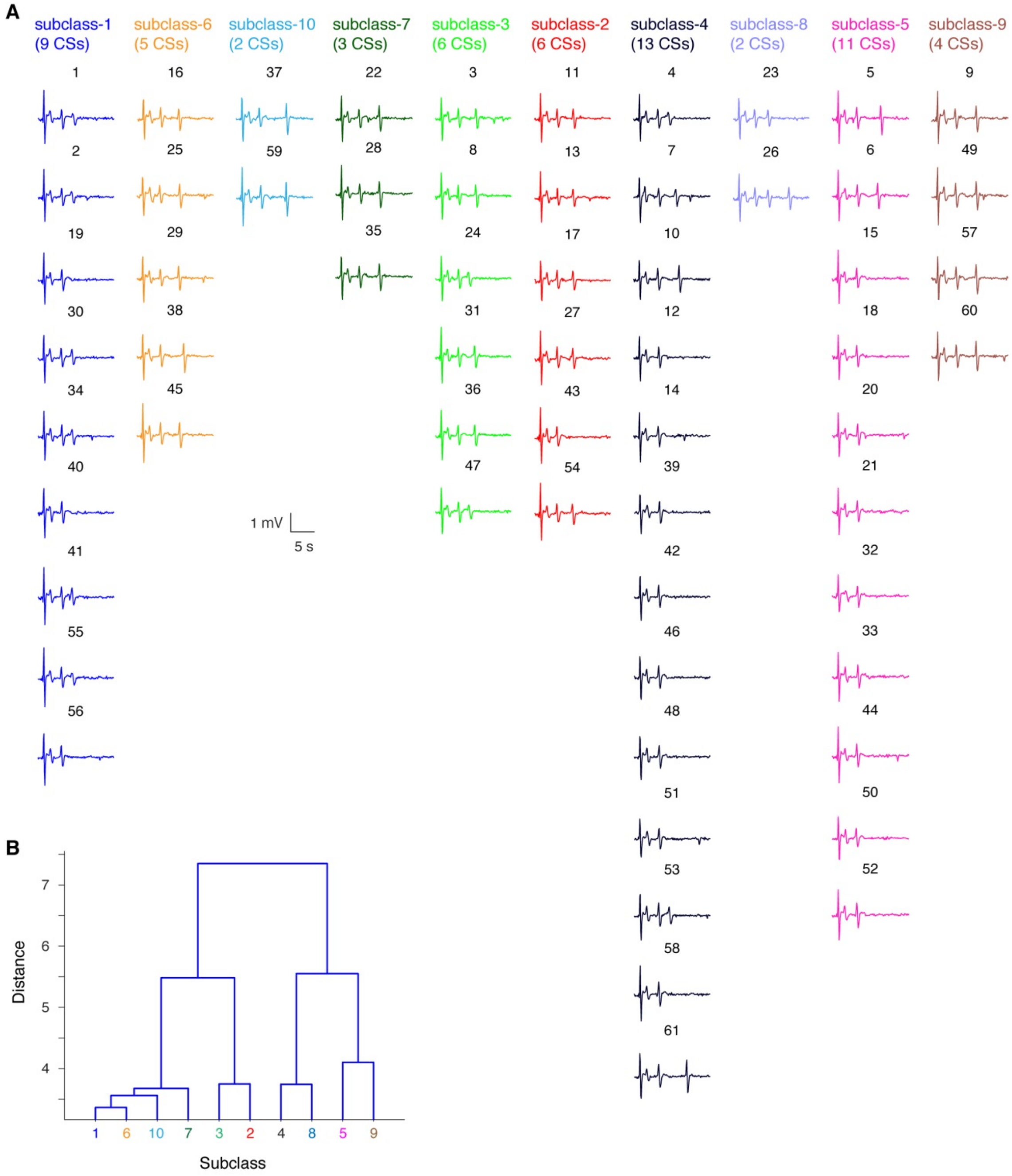

**Fig. S6. Classification of CS waveforms.** (A) Sixty-one CSs detected in a single record were classified into 10 subclasses according to hierarchical clustering analysis. The number of CSs in each subclass is shown in parentheses. Each of the 61 CS waveforms is color-coded depending on the classification (10 subclasses). (B) Dendrogram of the hierarchical clustering.

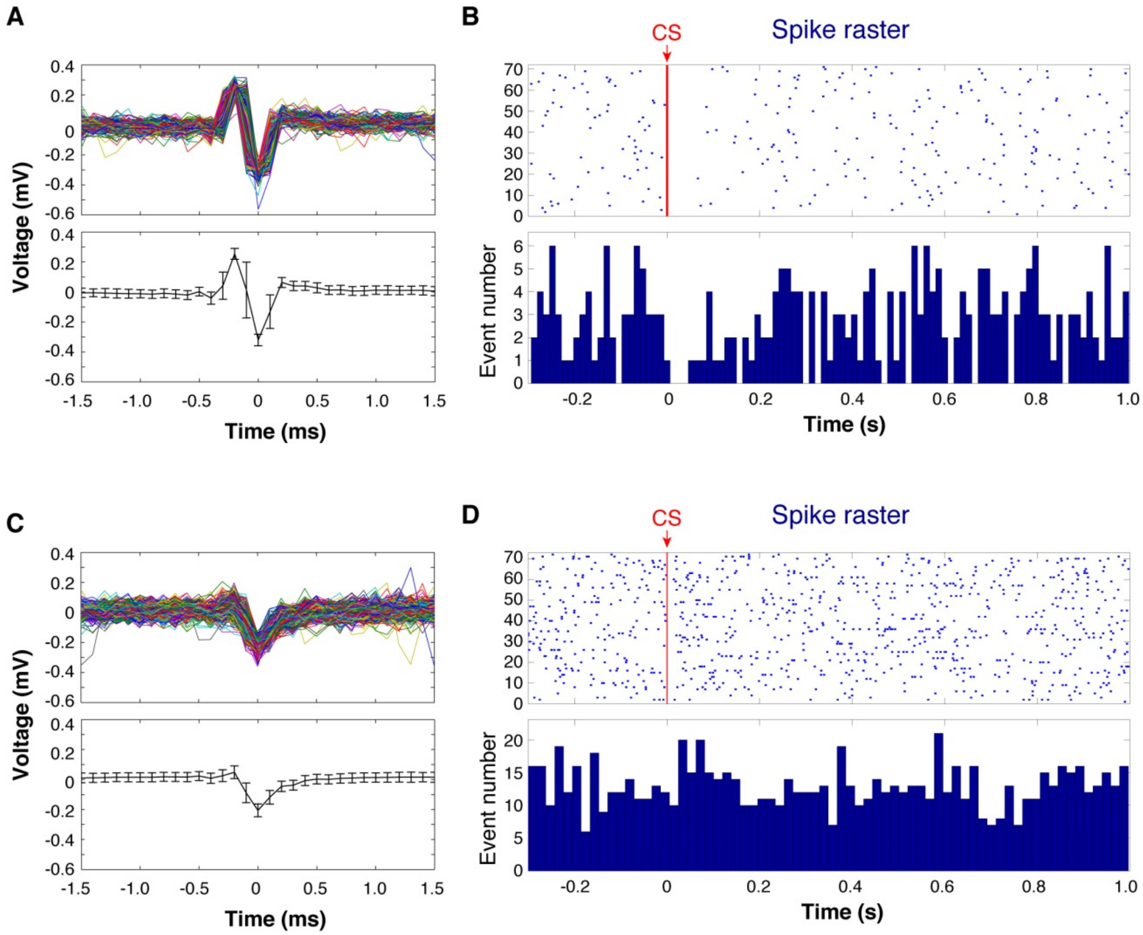


**Fig. S7. Spikes showing and not showing CS pauses.** (A and B) Spikes showing CS pauses. (A) The spikes observed in the data shown in Fig. 5. Waveforms of 318 spikes detected from a continuous 730-s record are superimposed (top panel). The average waveform is shown in the bottom panel. Error bars correspond to standard deviations. (B) Raster plot of the spikes shown in (A) aligned to CSs indicated by a red vertical line (top panel). Post-CS time histogram (lower panel). (C and D) Spikes not showing CS pauses. (C) Small amplitude spikes observed in the data shown in Fig. 5. Waveforms of 928 spikes detected from a continuous 730-s record are superimposed (top panel). The average waveform is shown in the bottom panel. Error bars correspond to standard deviations. (D) Raster plot of the spikes shown in (C) aligned to CSs indicated by a red vertical line (top panel). Post-CS time histogram (lower panel).
